# Genetic diversity of tilapia lake virus genome segment 1 from 2011 to 2019 and a newly validated semi-nested RT-PCR method

**DOI:** 10.1101/848903

**Authors:** Suwimon Taengphu, Pakkakul Sangsuriya, Kornsunee Phiwsaiya, Partho Pratim Debnath, Jerome Delamare-Deboutteville, Chadag Vishnumurthy Mohan, Ha Thanh Dong, Saengchan Senapin

## Abstract

The gene of RNA viruses, encoding RNA-directed RNA polymerase (RdRp) is relatively conserved due to its crucial function in viral genome replication and transcription making it a useful target for genetic diversity study and PCR detection. In this study, we investigated the genetic diversity of 21 tilapia lake virus (TiLV) genome segment 1 sequences predictively coding for RdRp subunit P1. Those sequences were obtained from infected fish samples collected in Ecuador, Israel, Peru, and Thailand between 2011 and 2019 (nine sequences from this study and 12 sequences from GenBank). Primers were then designed from the highly conserved regions among all 21 TiLV segment 1 sequences and used in semi-nested RT-PCR condition optimization. The result revealed that all 21 TiLV segment 1 sequences showed 95.00-99.94 and 99.00-100% nucleotide and amino acid sequence identity, respectively. These isolates were phylogenically clustered into three separate genetic clades, called i) Israeli-2011 clade (containing of TiLV isolates from Israel collected in 2011, Ecuador, and Peru isolates), ii) monophyletic Israel-2012 clade (containing only TiLV isolates collected from Israel in 2012), and iii) Thai clade (containing only sequences obtained from Thailand isolates). The newly established PCR protocol was 100 times more sensitive than our previous segment 3-based protocol when comparatively assayed with RNA extracted from infected fish. The assay was also shown to be specific when tested against negative control samples, i.e. RNA extracted from clinical healthy tilapia and from bacterial and viral pathogens (other than TiLV) commonly found in aquatic animals. Validation experiment with RNA extracted from naturally infected fish specimens collected in 2013-2019 yielded positive test results for all samples tested, confirming that our newly designed primers and detection protocol against TiLV segment 1, have a potential application for detection of all current genetic variants of TiLV.

## Introduction

A novel disease termed syncytial hepatitis of tilapia (SHT) associated with high mortality in farmed tilapia (Ferguson et al. 2014) caused by Tilapia lake virus (TiLV) (Eyngor et al. 2014). TiLV is a segmented RNA virus that has been taxonomically assigned to *Tilapia Tilapinevirus* (Bacharach et al. 2016; Adams et al. 2017). TiLV has been reported in 16 tilapia farming countries and this is likely to increase due to some underreporting (Jansen et al. 2018; Pulido et al. 2019). Rapid and accurate detection of the virus is crucial for selection of TiLV-free fish broodstock in view of the vertical transmission possibilities of the virus from parents to offspring (Yamkasem et al. 2019; Dong et al. 2020) and to prevent disease spread through movement of live fish for aquaculture (Dong et al. 2017a; Jansen et al. 2018). Following the characterization of the TiLV genomes (Eyngor et al. 2014; Bacharach et al. 2016), several PCR methods have been published, including RT-PCR (Eyngor et al. 2014), nested RT-PCR (Kembou Tsofack et al. 2017), semi-nested RT-PCR (Dong et al. 2017b), RT-qPCR (Kembou Tsofack et al. 2017; Tattiyapong et al.2018; Waiyamitra et al. 2018), and RT-LAMP (Phusantisampan et al. 2019; Yin et al. 2019). Most of the aforementioned methods used genome segment 3 of TiLV as the target for primer design. There are insufficient information and understanding on genetic diversity and functional role of segment 3. Due to the nature of RNA viruses, we hypothesized that there was a certain level of sequence variation in the TiLV genomes. If so, there might be a possibility for mismatches of the designed primers and thus the current molecular detection methods may not be applicable for all genetic variants, if any, of TiLV.

So far, the functions of the putative protein products of the 10 genomic segments of TiLV have not been elucidated, except for the sequence of segment 1, that has weak percent identity (~17% amino acid identity, 37% sequence coverage) to the *PB1* gene of influenza C virus (Bacharach et al. 2016). Influenza virus *PB1* gene encodes for RNA-directed RNA polymerase (RdRp) catalytic subunit sometimes called polymerase basic protein 1 or RNA-directed RNA polymerase subunit P1. Generally, sequences encoding the *RdRp* genes of RNA viruses show a reasonably high level of conservation due to their crucial replication and transcription functions. *RdRp* gene was therefore considered a useful molecular marker for virus classification and detection (Koonin and Dolja 1993; Culley et al. 2003; Culley and Steward 2007; Senapin et al. 2007). Therefore, this study aims to i) investigate the genetic diversity of TiLV genome segment 1 from TiLV-infected fish samples and sequences available from GenBank and ii) to develop a new semi-nested RT-PCR method based on primers designed from highly conserved regions of TiLV genome segment 1 for disease diagnosis and surveillance.

## Materials and methods

### Fish samples and RNA preparation

For amplification of TiLV genome segment 1, nine TiLV-infected specimens obtained from years 2013 to 2019 were used. They were archived specimens from our previous works (Dong et al.2017a; Dong et al. 2017b) and a newly collected set (Table 1). Suspected TiLV-infected fish and experimentally challenged fish samples (Table 2) were subjected to TiLV diagnosis using the newly developed semi-nested RT-PCR described below. RNA was extracted from the fish tissues using Trizol reagent (Invitrogen) following protocols recommended by the manufacturer. Quality and quantity of the obtained RNA was measured by spectrophotometry at 260 and 280 nm. RNA extracted from clinically healthy red and Nile tilapia tested negative for TiLV were used in negative control PCR reactions.

**Table 1.** Sources of TiLV sequences used to create the consensus segment 1 sequences in this study

| No. | Code | Sample collection | Accession no. | Fish host | Fish health status | Ref. |
| --- | --- | --- | --- | --- | --- | --- |
| 1 | TH-2013 | Jan-2013 | MN687685 | Nile tilapia fertilized eggs | Not specified | This study |
| 2 | TH-2014 | Aug-2014 | MN687695 | Nile tilapia fry | 90% mortality | This study |
| 3 | TH-2015 | Aug-2015 | MN687705 | Nile tilapia fingerling | 50% mortality | This study |
| 4 | TH-2016-CN | Dec-2016 | MN687725 | Red tilapia juvenile | 90% mortality | This study |
| 5 | TH-2016-CU | Dec-2016 | MN687715 | Tilapia fingerling | 20% mortality | This study |
| 6 | TH-2017 | 2017 | MN687735 | Nile tilapia | 90% mortality | This study |
| 7 | TH-2018-K | Aug-2018 | MN687755 | Nile tilapia juvenile | 50% mortality | This study |
| 8 | TH-2018-N | Jul-2018 | MN687745 | Red tilapia fingerling | 40-60% mortality | This study |
| 9 | TH-2019 | Feb-2019 | MN687765 | Nile tilapia fingerling | 30-50% mortality | This study |
| 10 | TH-2016-TV1 | Feb-2016 | KX631921 | Red tilapia | 20-90% mortality | Surachetpong et al. 2017 |
| 11 | TH-2015-TV2 | Dec-2015 | KX631931 | Red tilapia |  |  |
| 12 | TH-2016-TV3 | Jan-2016 | KX631932 | Red tilapia |  |  |
| 13 | TH-2016-TV4 | Jan-2016 | KX631933 | Nile tilapia |  |  |
| 14 | TH-2016-TV5 | Jan-2016 | KX631934 | Red tilapia |  |  |
| 15 | TH-2016-TV6 | May-2016 | KX631935 | Red tilapia |  |  |
| 16 | TH-2016-TV7 | May-2016 | KX631936 | Nile tilapia |  |  |
| 17 | TH-2018-WVL18053-01A | Apr-2018 | MH319378 | Nile tilapia | Not specified | NCBI |
| 18 | IL-2011- Til-4-2011 | 01-May-2011 | KU751814 | Tilapia | >80% mortality | Eyngor et al. 2014; Bacharach et al. 2016 |
| 19 | IL-2012-AD-2016 | 14-Aug-2012 | KU552131 | Hybrid tilapia | Not specified | NCBI |
| 20 | EC-2012 | 2012 | MK392372 | Nile tilapia | Up to 90% mortality | Subramaniam et al. 2019 |
| 21 | PE-2018-F3-4 | Feb-2016 | MK425010 | Nile tilapia | apparently healthy | Pulido et al. 2019 |

Country code: TH, Thailand, IT, Israel, EC, Ecuador, PE, Peru
Red tilapia (*Oreochromis* spp.); Nile tilapia (*Oreochromis niloticus*); Hybrid tilapia (*Oreochromis niloticus* x *Oreochromis aureus*)

**Table 2.** Sample sets subjected to TiLV detection using the present semi-nested RT-PCR segment 1 protocol

|  | Fish stage | Type of sample | Year collected | No. of +ve/tested sample |
| --- | --- | --- | --- | --- |
| Set 1 | Nile and hybrid red tilapia, various stages | Archived extracted RNA from clinically sick fish | 2013-2019 | 9/9 |
| Set 2 | Nile tilapia, broodstock and eggs | Experimentally TiLV infected samples | 2019 | 19/20* |
| Set 3 | Hybrid red tilapia, fingerlings | Frozen specimens of clinically sick fish | 2019 | 4/4 |
|  |  | Frozen specimens of clinically healthy fish | 2019 | 4/4 |
| Set 4 | Nile tilapia, broodstock | Blood from apparently healthy fish preserved in Trizol | 2019 | 0/8 |
| Set 5 | Nile tilapia, broodstock | Blood from apparently healthy fish preserved in Trizol | 2019 | 0/6 |
| Set 6 | Nile tilapia, fingerlings | Frozen specimens of clinically healthy fish | 2019 | 0/3 |
| Set 7 | Nile tilapia, broodstock | Blood from apparently healthy fish preserved in Trizol | 2019 | 0/40 |
|  |  | Liver biopsy sample preserved in Trizol | 2019 | 0/40 |
| Total |  |  |  | 36/134 |

\*13/20 samples tested positive by TiLV segment 3 protocol

### Amplification of TiLV genomic segment 1

Primers used for amplification of the putative open reading frame (ORF) of TiLV genomic segment 1 were TiLV-S1-F; 5’-ATG TGG GCA TTT CAA GAA-3’ and TiLV-S1-R; 5’-TTA GCA CCC AGC GGT GGG CT-3’ as previously described by Pulido et al. 2019. They were designed based on the segment 1 sequence of the TiLV Israeli isolate TiL-4-2011 (accession number KU751814, Eyngor et al, 2014, Bacharach et al, 2016). RNA extracted from 9 TiLV-infected fish specimens obtained as described above were individually used as template. RT-PCR reaction of 25 μl composed of 200 ng RNA template, 300 nM of each primer, 1 μl of SuperScript III RT/Platinum Taq Mix (Invitrogen), and 1x supplied buffer (containing 0.2 mM of each dNTP, 2.6 mM MgSO4, and stabilizers). Amplification profiles consisted of a reverse transcription step at 50 °C for 30 min; 35 cycles of 94 °C for 30 s, 55 °C for 30 s, and 72 °C for 1.5 min; and a final extension step at 72 °C for 5 min. After agarose gel electrophoresis, expected 1,560 bp amplicons obtained from each of the nine reactions were removed from the gel, purified and cloned into pGEM-T easy vector (Promega). Recombinant clones were then sent to Macrogen (South Korea) for DNA sequencing using T7 promoter and SP6 promoter primers.

### DNA sequence analysis

A total of 21 TiLV segment 1 sequences listed in Table 1 were used for comparison and phylogenetic relationship analysis. Nine sequences were obtained from the present study and other 12 sequences were retrieved from the GenBank database. Multiple sequence alignments were performed using MEGA 7 (Kumar et al. 2016) and a consensus sequence of 1,559 nucleotide residues out of the putative 1,560 nucleotide-ORF segment 1 were obtained. Deduced amino sequences of 519 nucleotide residues were translated from these consensus TiLV segment 1 using ExPASy translate tool and then used for sequence comparison. Putative PB domain of the translated sequences was predicted using InterPro database (https://www.ebi.ac.uk/interpro/). Phylogenetic tree based on TiLV segment 1 nucleotide sequences was constructed using Maximum-Likelihood with GTR+G (General Time Reversible model + Gamma distributed) method as suggested by a best model feature of the MEGA 7 program. An ORF coding for a putative *PB1* of influenza virus C isolate Ann Arbor 1950 (NC_006308) was used as outgroup. Pairwise distance analysis was also conducted using MEGA 7.

### Development of a new semi-nested RT-PCR method based on segment 1

In this study, a new semi-nested RT-PCR detection of TiLV was developed. Primers were designed from the highly conserved regions from the multiple sequence alignments of the 21 sequences of TiLV segment 1. These sequences derived from TiLV isolates collected from the period from 2011 to 2019 (Table 1). Primer specificity was initially confirmed *in silico* with the NCBI primer-blast tool. Primers TiLV/nSeg1F; 5’-TCT GAT CTA TAG TGT CTG GGC C-3’ and TiLV/nSeg1R; 5’-AGT CAT GCT CGC TTA CAT GGT-3’ with an expected amplified product of 620 bp were used in the first round RT-PCR. Primers TiLV/nSeg1F and TiLV/nSeg1RN; 5’-CCA CTT GTG ACT CTG AAA CAG-3’ with an expected product of 274 bp were employed in the second round PCR. Amplification conditions such as annealing temperature, number of cycles, buffer and primer concentration were evaluated to obtain optimized conditions. Consequently, the first RT-PCR reaction of 25 μl composed of 200 ng of RNA template, 400 nM of each primer, 0.5 μl of SuperScript III RT/Platinum Taq Mix (Invitrogen), and 1x of supplied buffer. Amplification profiles consisted of a reverse transcription step at 50 °C for 30 min; a denaturation step at 94 °C for 2 min, 30 PCR cycles of 94 °C for 30 s, 60 °C for 30 s, and 72 °C for 30 s; and a final extension step at 72 °C for 2 min. Positive control consisted of a reaction containing RNA extracted from TiLV-infected red tilapia as template while no template added in the negative control. 5 μl of product from the first round PCR was then used as template in the second round PCR reaction of 25 μl containing 500 nM of primer TiLV/nSeg1F, 600 nM of primer TiLV/nSeg1RN, 0.16 mM of each dNTP, 0.8 mM MgCl2, 1 unit of Platinum Taq DNA polymerase (Invitrogen), and 1.2x supplied buffer. Thermocycling conditions consisted of a 5 min initial denaturation step at 94 °C followed by 30 thermocycles and a final extension step described above. 10 μl of the amplified products were analyzed by 1.5% agarose gel electrophoresis stained with ethidium bromide or RedSafe DNA staining dye.

### Validation of TiLV segment 1 semi-nested RT-PCR assay

Detection specificity of the assay was performed with RNA extracted from common viral and bacterial pathogens found in aquatic animals as listed in Table 3. Viral infected fish tissues were subjected to RNA extraction using Trizol reagent as mentioned above. For bacterial RNA isolation, each bacterial strain was cultured in appropriated broth medium of 5 mL overnight at 30 °C with shaking at 200 rpm. After centrifugation, bacterial cell pellets were washed once with nuclease-free water and homogenized with Trizol reagent followed by the manufacturer’s protocol. Detection sensitivity of the TiLV segment 1 protocol (this study) was performed by comparison with our previous semi-nested PCR protocol targeting TiLV segment 3 (Dong et al. 2017b). This was done using serially diluted RNA template extracted from 3 individual TiLV-infected fish. Two fish samples were assayed by Centex Shrimp in Thailand while another by WorldFish Khulna laboratory, Bangladesh. The newly established semi-nested RT-PCR protocol was then employed to screen for TiLV infection in both clinically healthy and clinically sick fish samples, including representative from archived samples as well as naturally and experimentally infected fish (Tables 1 and 2).

**Table 3.**
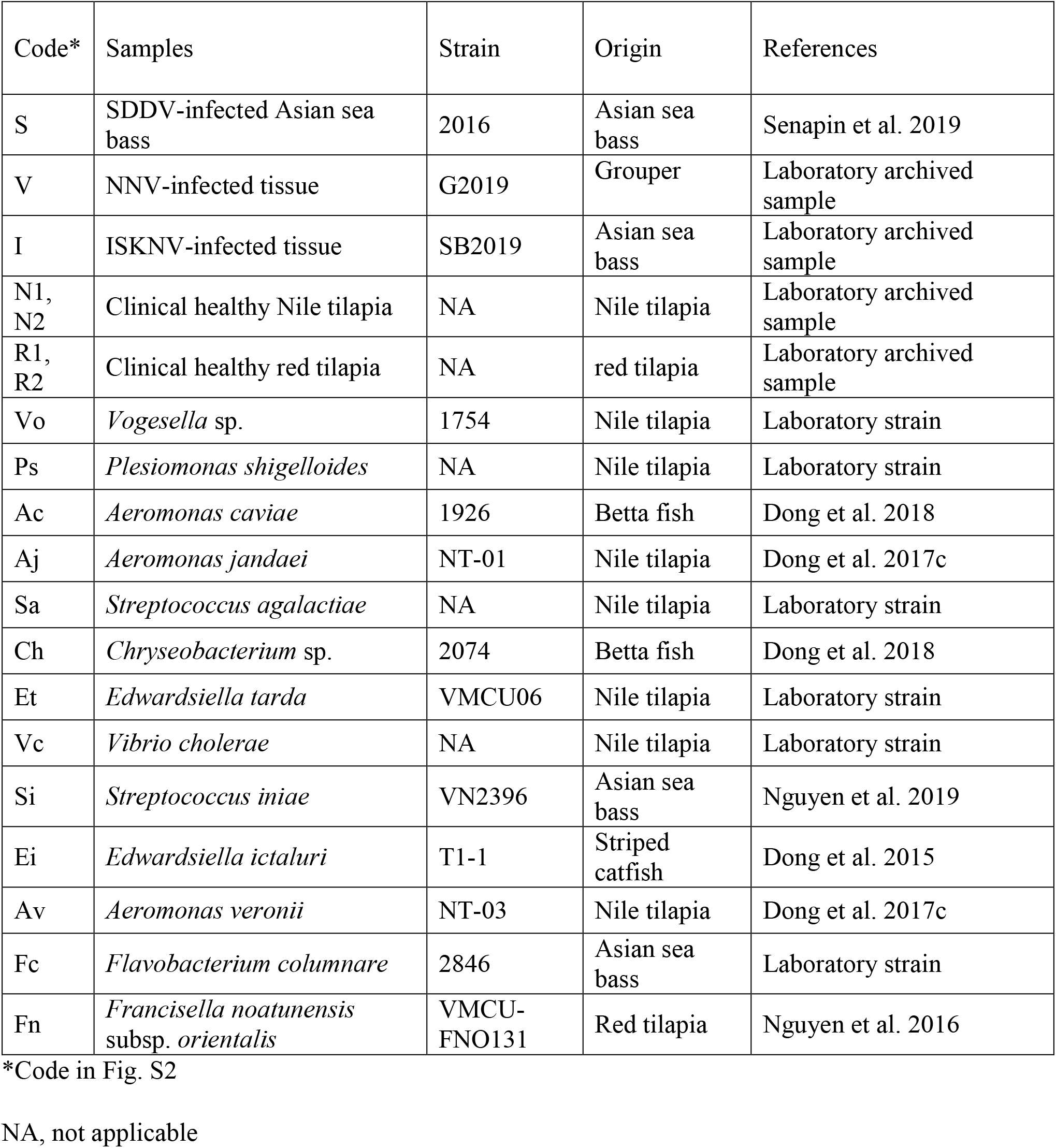
RNA samples used for specificity assay in this study

| Code* | Samples | Strain | Origin | References |
| --- | --- | --- | --- | --- |
| S | SDDV-infected Asian sea bass | 2016 | Asian sea bass | Senapin et al. 2019 |
| V | NNV-infected tissue | G2019 | Grouper | Laboratory archived sample |
| I | ISKNV-infected tissue | SB2019 | Asian sea bass | Laboratory archived sample |
| N1, N2 | Clinical healthy Nile tilapia | NA | Nile tilapia | Laboratory archived sample |
| R1, R2 | Clinical healthy red tilapia | NA | red tilapia | Laboratory archived sample |
| Vo | *Vogesella* sp. | 1754 | Nile tilapia | Laboratory strain |
| Ps | *Plesiomonas shigelloides* | NA | Nile tilapia | Laboratory strain |
| Ac | *Aeromonas caviae* | 1926 | Betta fish | Dong et al. 2018 |
| Aj | *Aeromonas jandaei* | NT-01 | Nile tilapia | Dong et al. 2017c |
| Sa | *Streptococcus agalactiae* | NA | Nile tilapia | Laboratory strain |
| Ch | *Chryseobacterium* sp. | 2074 | Betta fish | Dong et al. 2018 |
| Et | *Edwardsiella tarda* | VMCU06 | Nile tilapia | Laboratory strain |
| Vc | *Vibrio cholerae* | NA | Nile tilapia | Laboratory strain |
| Si | *Streptococcus iniae* | VN2396 | Asian sea bass | Nguyen et al. 2019 |
| Ei | *Edwardsiella ictaluri* | T1-1 | Striped catfish | Dong et al. 2015 |
| Av | *Aeromonas veronii* | NT-03 | Nile tilapia | Dong et al. 2017c |
| Fc | *Flavobacterium columnare* | 2846 | Asian sea bass | Laboratory strain |
| Fn | *Francisella noatunensis* subsp. *orientalis* | VMCU-FNO131 | Red tilapia | Nguyen et al. 2016 |

\*Code in Fig. S2
NA, not applicable

## Results

### Genome segment 1 based phylogenetic relationship of TiLV isolates

After amplification of TiLV genome segment 1 from the nine TiLV-infected tilapia, their sequences were compared to that of 12 sequences retrieved from the GenBank database. Consequently, 21 TiLV segment 1 sequences obtained from samples collected in 2011 to 2019 were used for phylogenetic analysis. The results showed that the TiLV segment 1 sequences were clustered into three separate clades namely Israeli-2011 clade, Israeli-2012 clade, and Thai clade while the *RdRp PB1* sequence of influenza C virus used as an outgroup diverged into another linage (Fig. 1). The Thai clade comprised all studied isolates from Thailand was closely related to the monophyletic Israel-2012 clade, whereas Israeli-2011 clade composed of three TiLV isolates (i.e. IL-2011-Til-2011, EC-2012, PE-2018-F3-4) from Israel, Ecuador, and Peru respectively. Additionally, the phylogenetic tree revealed 2 subclades within the Thai clade (Fig. 1). Further investigation using pairwise distance analyses, revealed that the three proposed clades diverged among each other from 0.029 to 0.044 (i.e. 2.9-4.4%), with the lowest and highest values being between Israeli-2012 & Thai clades and Israeli-2011 & Thai clades, respectively. Average divergence within Israeli-2011 clade and within Thai clade were 0.033 and 0.035 (i.e. 3.3-3.5%), respectively. The pairwise genetic distance is shown in Table S1.

**Figure 1.**
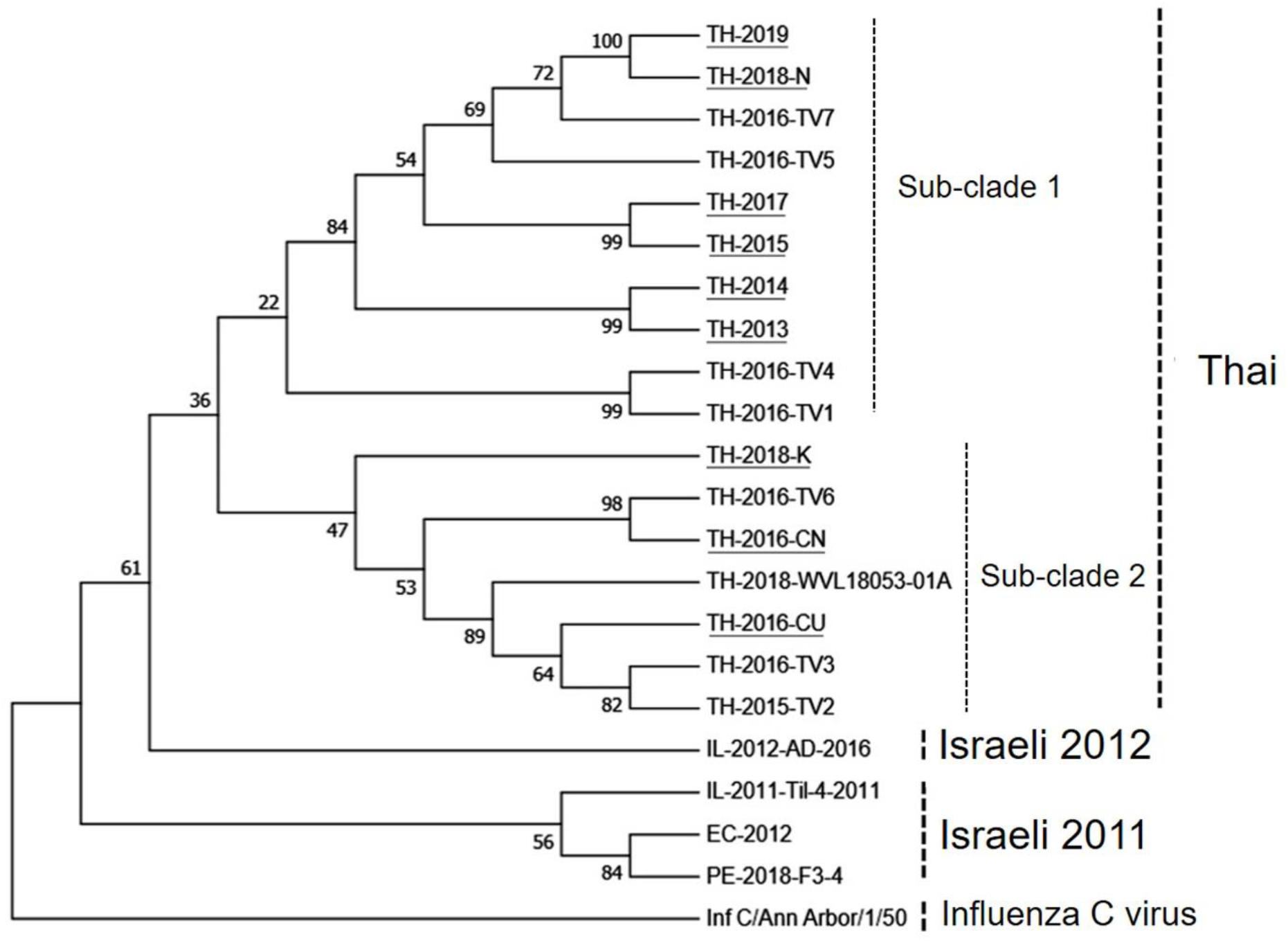
Maximum-likelihood phylogenetic trees based on TiLV genomic segment 1 *PB1* nucleotide sequences depicting TiLV being clustered into 3 clades. Isolates with and without underlines represent sequences obtained from this study and those from the GenBank database, respectively. Codes of the viral isolates are listed in Table 1. The influenza C virus sequence was used as outgroup. Percent bootstrap values from 1,000 replications are shown.

### Conserved regions of TiLV segment 1 gene sequences and primer design

Segment 1 gene sequences of the 21 TiLV isolates were compared using Clustal Omega and MEGA 7 programs. Among the 21 sequences, there were 94.29-99.94 and 97.11-100% identity at the nucleotide and amino acid sequence levels, respectively. Noted that among the 17 Thai isolates, the respective percentages were 95.00-99.94 and 97.50-100%. PB domains and four RdRp motifs were detected from all of the deduced amino acid sequences (Fig. S1). Based on prediction published by Chu et al. 2012 for influenza virus and Bacharach et al. 2016 for TiLV, the four RdRp motifs of TiLV containing motif I ^48^**<u>TGD</u>**L^51^, motif II ^260^CPG**<u>GM</u>**L**<u>MGMF</u>**^269^, motif III ^282^DRFLSF**<u>SDDF</u>**ITS^294^, and motif IV ^312^CH**<u>N</u>**L**<u>S</u>**L**<u>KKSYI</u>**^322^; where bold and underlined letters represent residues matching to those of influenza virus sequences. Those specific residues were found in all 21 TiLV isolates (Fig. S1).

The obtained multiple nucleotide sequence alignments displayed conserved regions along segment 1 gene sequences for the 21 TiLV isolates (Fig. 2). In this study, we aimed to design conserved primers to be used in a semi-nested RT-PCR assay enabling amplification of all genetic variability (known to date) of TiLV isolates if any. Primers TiLV/nSeg1F, TiLV/nSeg1R, and TiLV/nSeg1RN are 21-22 nucleotides long and locate at nucleotide positions 717-737, 1315-1336, and 1063-1083; all locate in the segment 1 ORF, showing 100% identity among all of the 21 TiLV isolates analyzed (Fig. 2). Amplicon sizes of 620 and 274 bp were expected to be produced from primers TiLV/nSeg1F and TiLV/nSeg1R; and TiLV/nSeg1F and TiLV/nSeg1RN, respectively (Fig. 2). Additionally, these primer pairs were shown, *in silico*, to be TiLV specific using primer-blast tool (https://www.ncbi.nlm.nih.gov/tools/primer-blast/). Thus, PCR condition optimization for TiLV detection using the newly designed primers was conducted as described below.

**Figure 2.**
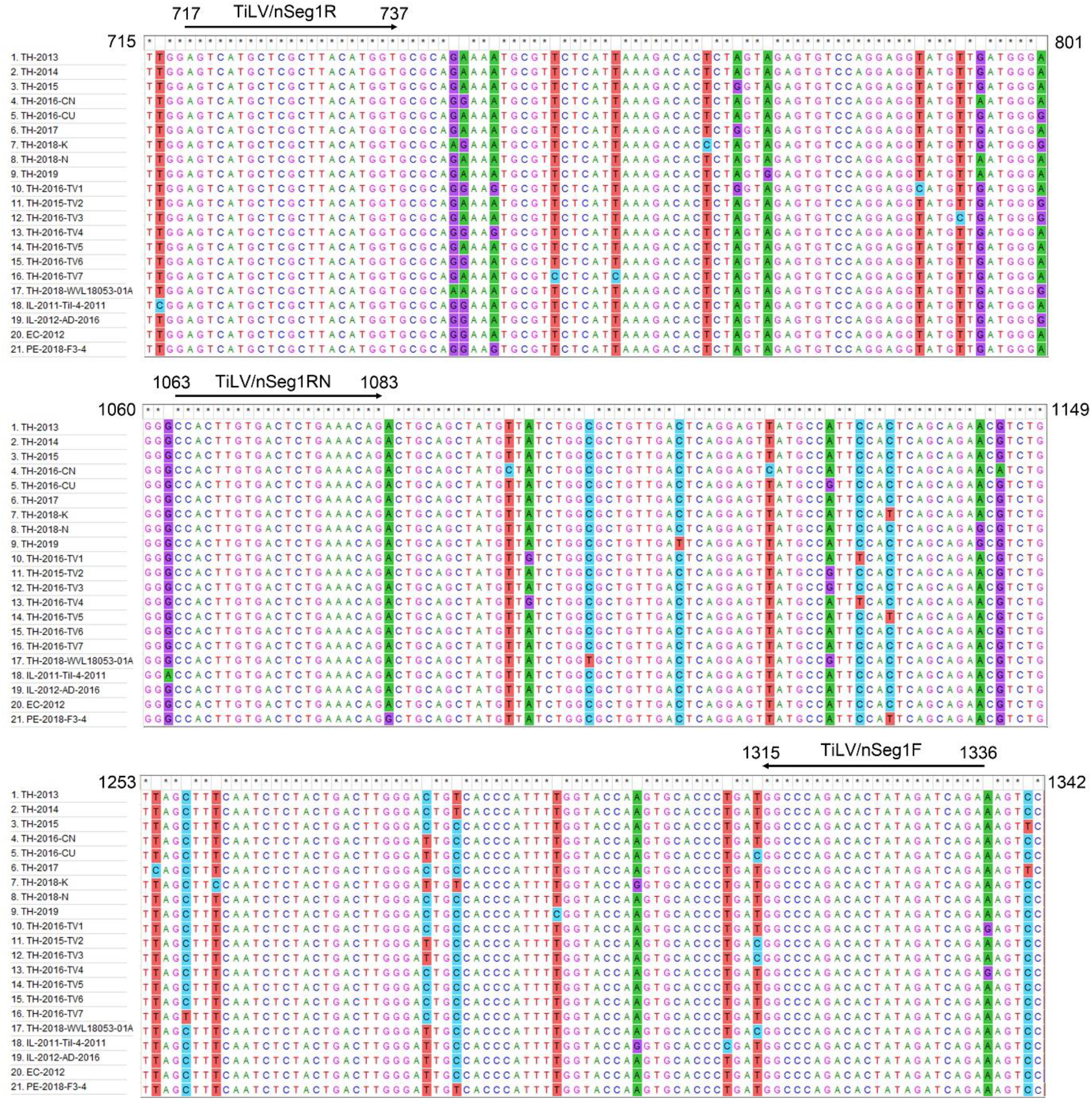
Multiple nucleotide sequence alignments of TiLV segment 1 *PB1* genes of 21 viral isolates. A partial portion of an alignment is shown. Numbers represent nucleotide positions of the coding strand in the putative *PB1* ORF. Arrows indicate sites that were used to design primers for the semi-nested RT-PCR protocol developed in this study. * marks nucleotide identity. Columns with differences in colors indicate nucleotide variability.

### Development of a validated TiLV detection protocol

The designed primer pairs described above were evaluated for their optimal annealing temperatures which were found to be 60°C for both first and nested amplification (figure not shown). Other parameters including numbers of cycles, buffer and primer concentration were also evaluated and optimized. Finally, our new TiLV segment 1 detection protocol was shown to be specific for only TiLV when tested against other 16 viral and bacterial pathogens (Fig. S2). The detection sensitivity assay was conducted by comparing between the present newly established TiLV segment 1 based protocol and our previous method which was based on segment 3 (Dong et al. 2017b) using serially diluted RNA from three individual TiLV-infected fish specimens. Results revealed that the new protocol described in this study was 100 times more sensitive than the previous one (Fig. 3a). This assay was done in two different laboratories i.e. 2 samples (i.e. fish 1 and 2) performed in Thailand and fish 3 in WorldFish Khulna laboratory, Bangladesh. Note the presence of a ~1.1 kb band predicted to be derived from cross hybridization of the amplified products was visible at higher concentrated templates (Fig. 3a). Moreover, one set of specimens from experimentally TiLV infected fish (Table 2, set 2, n=20) obtained from our previous study (Dong et al. 2020) was assayed using both segment 1 and segment 3 based detection protocols (Fig 3b), also confirming higher sensitivity of the current semi-nested PCR detection targeting TiLV segment 1 gene. This was evidenced by the numbers of positive tests obtained for segment 1 and segment 3 protocols, which were 19 and 13 out of 20, respectively (Table 2). Representative test results are shown in Fig. 3b. Comparison of TiLV infected fish samples collected from Thailand (2013-2019) and from Peru in 2018 (Tables 1 and 2) confirmed that the new detection protocol can be used to detect TiLV from all the samples tested representing Thai clade and Israeli-2011 clade in this study (Fig. 4). In addition, in one population, both clinically sick fish and apparently healthy fish were all tested positive for TiLV segment 1 (Table 2, set 3). Remaining sets (Table 2, sets 4-7) from field samples subjected to the test for TiLV were all negative.

**Figure 3.**
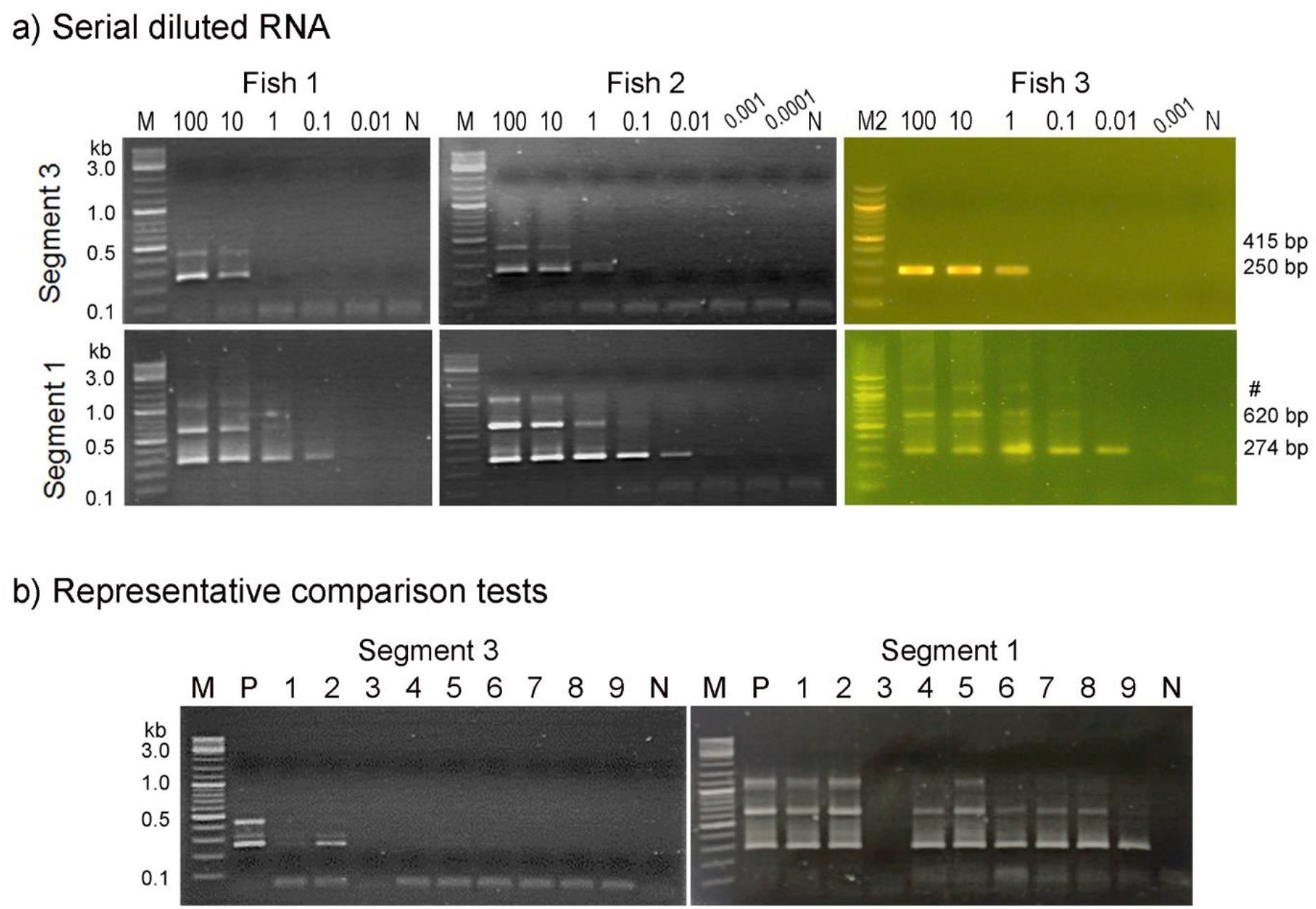
a) Sensitivity tests of two semi-nested RT-PCR protocols (segment 1 and 3) using serial dilutions of RNA from three TiLV-infected fish. A starting 100 ng of RNA from three individual TiLV-infected fish was 10-fold serially diluted and subjected to semi-nested RT-PCR protocols using primers targeting genome segment 3 (Dong et al. 2017b) and segment 1 (this study). M, 2 log DNA marker; N, no template control; #, cross hybridized PCR products. b) Representative comparison tests with 9 archived samples (1-9) obtained from Dong et al. (2020); P, positive control (infected tissue) and N, negative control (no RNA template).

**Figure 4.**
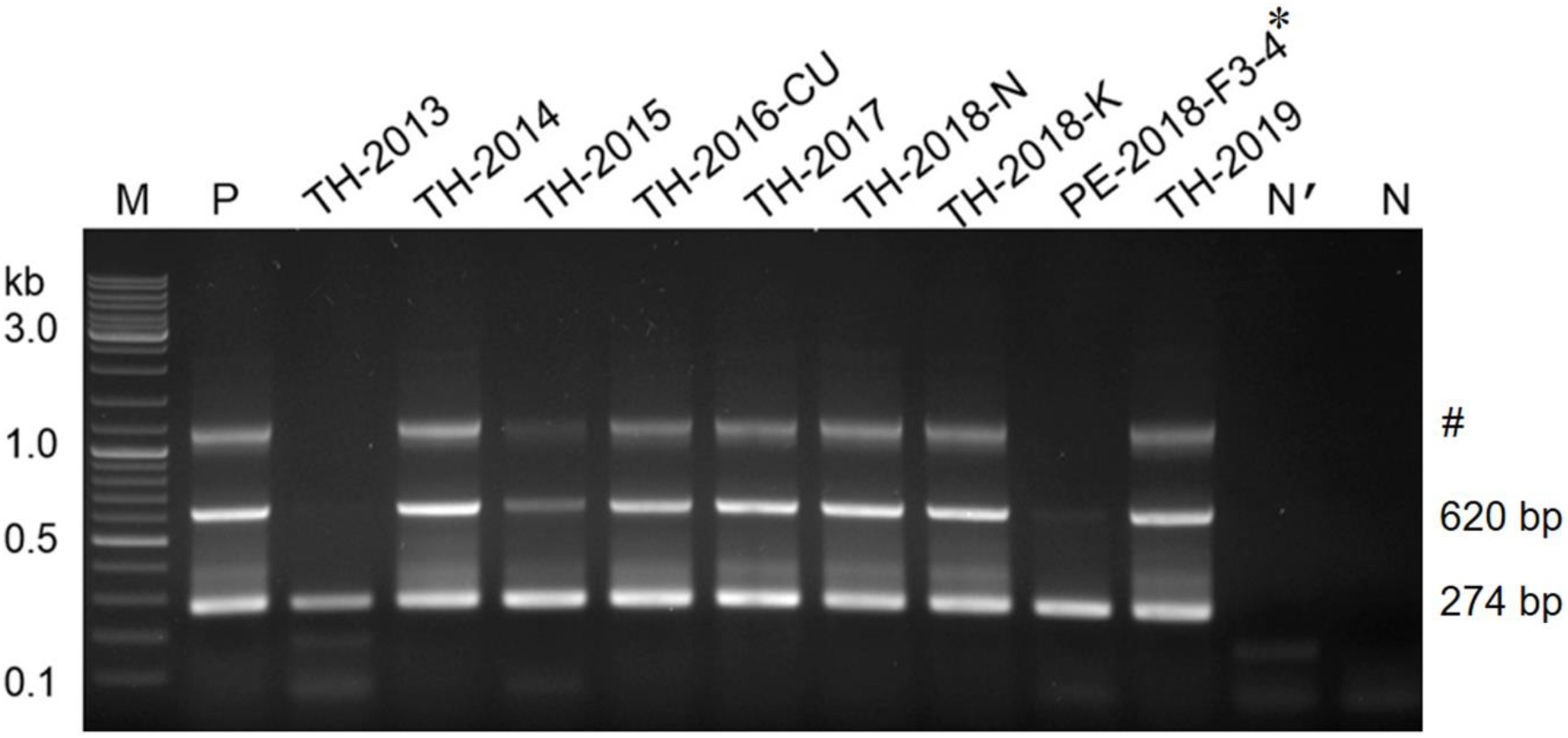
Validation of the newly developed semi-nested RT-PCR assays in detecting TiLV segment 1 from archived samples obtained from years 2013 to 2019. *sample from Peru described in Pulido et al. (2019); M, 2 log DNA marker; P, positive control (infected tissue); TH, samples from Thailand; PE, sample from Peru; N’, non-infected fish RNA; N, no template control; #, cross hybridized PCR products.

## Discussion

Recently, Pulido et al. 2019 proposed two genetic clades of TiLV (Israeli and Thai clades) based on multilocus sequence phylogenetic analysis (MLSA) of 8,305 nucleotides of 5 TiLV genomes. Due to limited number of complete TiLV genomes available in the GenBank database, this study, therefore, aimed to explore the genetic diversity of TiLV based on the open reading frame (ORF) of segment 1 *PB1* gene (1,560 nucleotides) from 21 TiLV isolates which comprised of the isolates from Israel, Peru, Ecuador, and Thailand. Although bootstrap values were not relatively high, phylogenetic tree inferred from the segment 1 dataset indicates three separate clades (Israeli-2011, Israeli-2012, and Thai clades). It is likely that the Israeli-2012 and Thai clades evolved from the Israeli-2011 clade. However, to gain a better understanding on TiLV evolution and how the virus emerged and widespread, larger number of complete TiLV genomes or sequences of segment 1 of TiLV from a broader origin should be made available. Note that no correlation between sequence variation or phylogenetic branches and disease severity was observed in this study.

Comparative nucleotide sequences revealed that TiLV segment 1 is relatively well conserved with 95.00-99.94% identity among all isolates used in this study, which is similar to that of Orthomyxoviruses such as Salmon isavirus RNA polymerase PB1 (97.60-99.86%) and Influenza C virus polymerase PB1 (97.12-99.34%) (BLAST search in this study). Taken together, the results from this study, suggest that TiLV segment 1 is a promising gene candidate for the study of the genetic diversity of TiLV globally.

Majority of tilapia farming countries are located in low and middle-income countries. In those countries, conventional PCR remains the preferable technique for many laboratories (in terms of costs and ease of use) compared to less accessible and more expensive quantitative PCR machine (Charoenwai et al. 2019). The semi-nested PCR method in this study therefore was designed based on conserved regions of genome segment 1 that was able to amplify representatives from both Thai, Israeli-2011 and Israeli-2012 clades. For its higher sensitivity and specificity, this new protocol, thus, might be able to reduce chance of false negative due to genetic variation if any.

Using serial dilutions of DNA plasmid containing insert of targeted TiLV segment 3 gene, our previous semi-nested RT-PCR targeting segment 3 (Dong et al. 2017b) has a detection limit of 7.5 copies per reaction. Similar approaches have been used in previously reported TiLV detection protocols (Kembou Tsofack et al. 2017; Tattiyapong et al. 2018). While detection limits are based on serial dilutions of DNA plasmid containing target gene, diagnostic assays in reality started from extracted RNA from infected fish, comprising both host and viral RNA. Thus, detection limit of the PCR assay for clinical samples might not be comparable with the results obtained from standard serial dilutions of DNA plasmid in the laboratory. This study, therefore, compared the sensitivity of two detection protocols together instead of using plasmid template to find out which protocol was the best for field detection purpose. Interestingly, inter-laboratory tests consistently proved that segment 1 PCR method was 100 times more sensitive than our previous segment 3 PCR protocol, when assayed with RNA extracted from diseased fish samples. In this study, segment 1 PCR detection method showed no cross-amplification against a panel of extracted RNA from various bacterial and viral pathogens of freshwater fish. When applied to field samples, this new segment 1 protocol was able to detect the virus not only from clinically sick fish but also from asymptomatically infected fish, indicating its potential use for screening early infected fish samples. Taken together, we recommend the use of this newly established segment 1 PCR method as an alternative tool for TiLV diagnosis and in active surveillance programs.

## Acknowledgments

This study was financially funded by a research grant from National Science and Technology Development Agency (NSTDA), Thailand (P-18-52583) and partially supported by the CGIAR Research Program on Fish Agri-Food Systems (FISH) led by WorldFish.

---

**Table S1.**
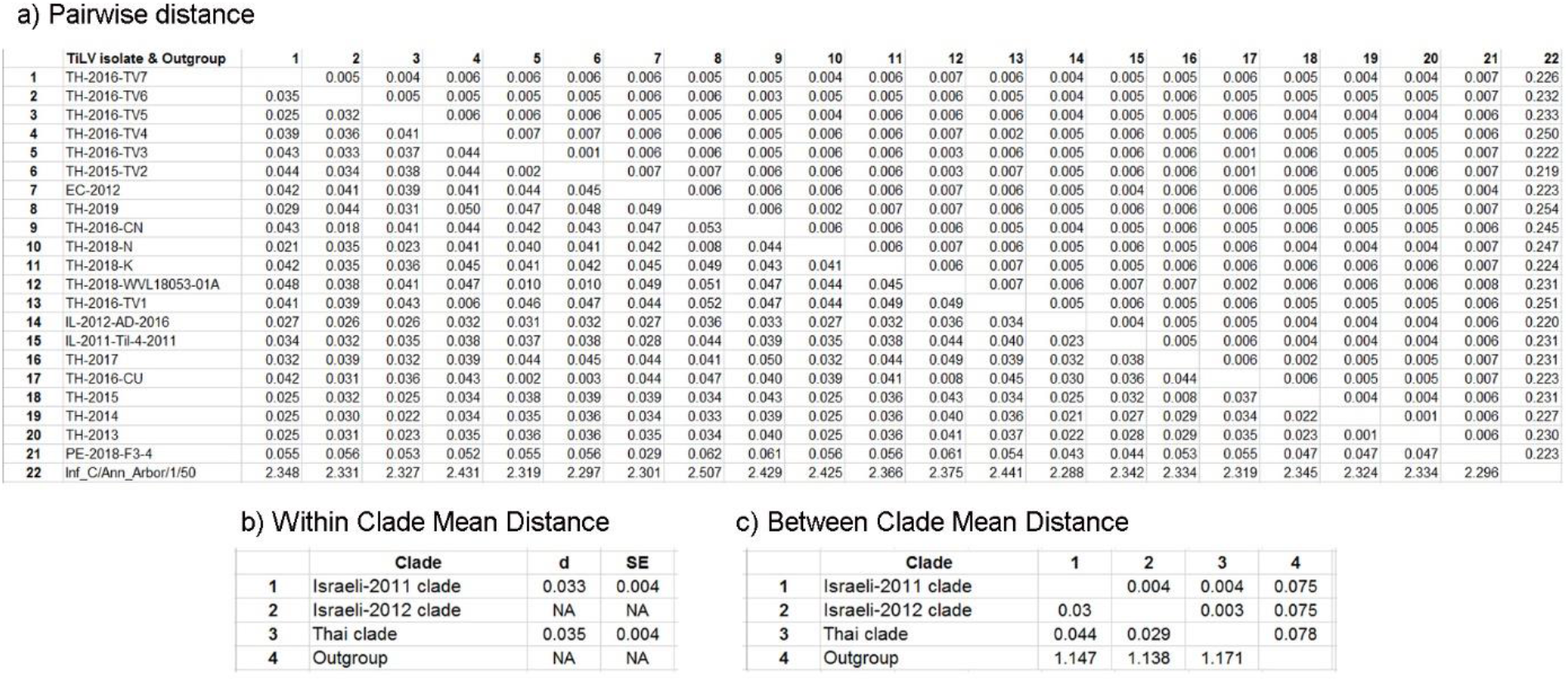
Pairwise genetic distance between TiLV isolates and influenza virus C as outgroup. (a) Divergence between sequences (b) Mean divergence within clades (c) Divergence between clades. The number of base substitutions per site from between sequences are shown below diagonal. Standard error estimate(s) are shown above the diagonal. Analyses were conducted from 1,000 bootstrap replicates using the Tajima-Nei model in MEGA 7 (Tajima and Nei 1984; Kumar et al.2016).

**Figure S1.**
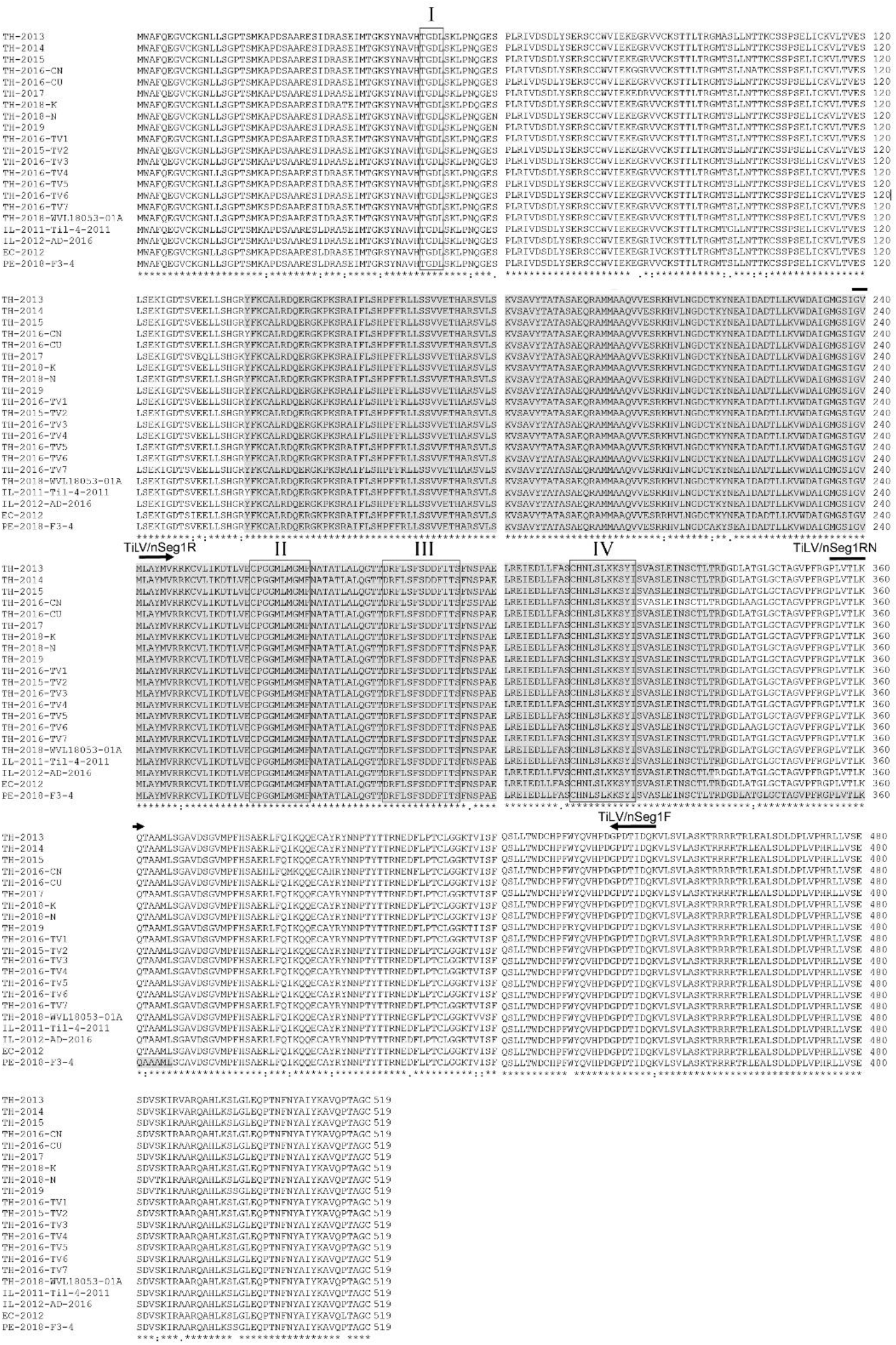
Multiple alignment of the deduced amino acid sequences of TiLV segment 1 PB1 coding genes of 21 viral isolates. Length of amino acid sequence is indicated on the right margin. *marks amino acid identity. The shaded amino acid sequences represent putative PB domains while the 4 predicted RdRp motifs are marked by boxes (I, II, II, and IV). Arrows indicate positions and orientation of the primers used in the newly developed TiLV detection assay.

**Figure S2.**
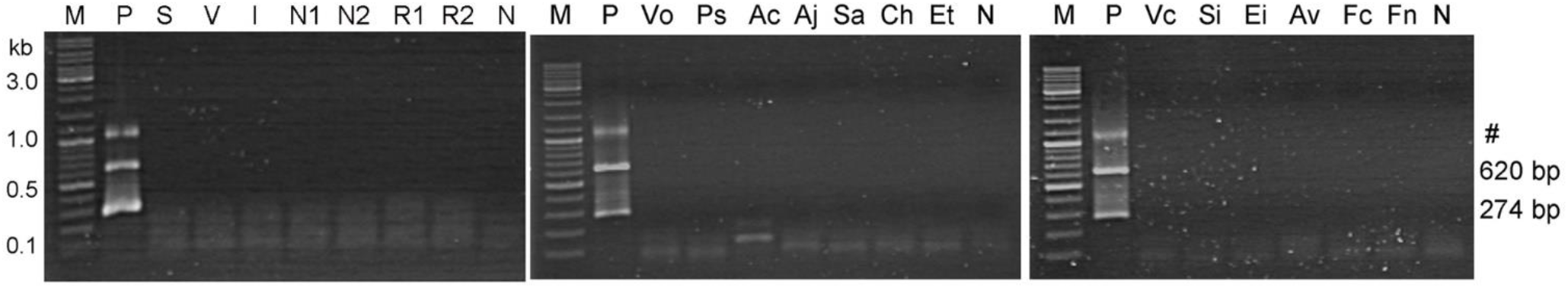
Specificity of the semi-nested TiLV detection assays. M, 2-log DNA marker (New England Biolabs); P, positive control using RNA extracted from TiLV-infected fish tissue as template; N, no template control; S, Scale Drop Disease Virus (SDDV); V, Nervous Necrosis Virus (NNV); I, Infectious Spleen and Kidney Necrosis Virus (ISKNV); N1/N2, clinically healthy Nile tilapia; R1/R2, clinically healthy red tilapia; Vo, *Vogesella* sp.; Ps, *Pleisiomonas shigelloides;* Ac, *Aeromonas caviae;* Aj, *Aeromonas jandaei;* Sa, *Streptococcus agalactiae;* Ch, *Chryseobacterium* sp.; Et, *Edwardsiella tarda;* Vc, *Vibrio cholerae;* Si, *Streptococcus iniae;* Ei, *E. ictaluri;* Av, *Aeromonas veronii*; Fc, *Flavobacterium columnare*; Fn, *Francisella noatunensis* subsp. *orientalis*. Details of bacterial isolates are shown in Table 2.

